# Dissecting Breast Cancer Heterogeneity Through Transcriptomics Insights of Diverse Etiological Factors for Common Biomarker Discovery

**DOI:** 10.1101/2024.10.20.619262

**Authors:** Mohammad Uzzal Hossain, Mariam Ahmed Mehak, SM Sajid Hasan, Mohammad Nazmus Sakib, A.B.Z. Naimur Rahman, Arittra Bhattacharjee, Zeshan Mahmud Chowdhury, Ishtiaque Ahammad, Md. Mehadi Hasan Sohag, Keshob Chandra Das, Md. Salimullah

**Author notes:** Joint first author (equally contributed to this work) (Mohammad Uzzal Hossain, Mariam Ahmed Mehak, SM Sajid Hasan). **Corresponding Author:** Dr. Md. Salimullah, Director General, Molecular Biotechnology Division, National Institute of Biotechnology, Ganakbari, Ashulia, Savar, Dhaka-1349, Bangladesh.

## Abstract

Breast cancer has many different causes, and the key to finding effective treatments is understanding the disease’s heterogeneity. The present study used three gene expression datasets from 110 female samples related to stress, drug and hormonal imbalance, diet and nutrition, and physical activity and light exposure at night to predict differential gene expression. Interestingly, all gene expression datasets shared 22 upregulated and 4 downregulated genes, regardless of etiology. This suggests these genes share the core molecular mechanism and the biological pathway that causes breast cancer. Notably, these genes were significantly enriched in some important pathways, including cycle regulation, endoplasmic reticulum stress, and transcriptional regulation, demonstrating their potential as therapeutic targets. Further, we found UBE2J2 from upregulated genes and ZCCHC7 from downregulated genes as the top hub and bottleneck genes, which may help network connectivity and functional gene interactions. Computational study further asserted the strong binding affinity of drug-target complexes. Later, molecular dynamics simulations confirmed the predicted drug-target complexes’ stability and dynamic behavior, demonstrating these two genes as potential therapeutic targets. The findings from this analysis provide the molecular basis into the complex interplay between diverse etiologic factors and breast cancer pathogenesis, paving the way for innovative biomarker-targeted therapies.

## Introduction

Breast cancer is a significant contributor to cancer-related morbidity and mortality globally, representing roughly 11.6% of all female malignancies and 6.6% of all cancer casualties. Breast cancer impacts 80% of those over the age of 50, with 40% being over 65. Epigenetic modifications and environmental factors drive this correlation. Strikingly, breast cancer is ranked as the second most diagnosed malignancy and the fifth leading cause of cancer-related mortality. The incidence rate is higher in developed countries, particularly in those with a high or very high Human Development Index (HDI)^1–4^. Consequently, comprehending the heterogeneity of this malignancy is of paramount importance. This heterogeneity is attributable to the several etiological variables that contribute to the development of breast cancer. Such factors include Excessive alcohol consumption which increases the risk of breast cancer, primarily due to elevated oestrogen levels and hormonal imbalances^5^. Excessive fat storage and increased Body Mass Index (BMI) levels significantly boost risks. Alcohol-induced nutritional deficiencies and both direct and indirect carcinogenic effects increase the likelihood of developing cancer. Alcohol consumption before pregnancy induces morphological alterations in breast tissue, making it vulnerable to carcinogenic events. Tobacco carcinogens in breast tissue increase the likelihood of mutations in oncogenes and tumour suppressor genes, including p53. Both active and passive smoking contribute to carcinogenic events, with a prolonged smoking history and genetic predisposition being key determinants^6,7^. Excessive consumption of saturated fats and ultra-processed meals, rich in sodium, fat, and sugar, predisposes persons to obesity, ultimately leads to breast cancer risk^6^. The natural circadian rhythm, which is affected by physical activity and nighttime light exposure (LAN), may be linked to breast cancer. This leads that nighttime exposure to artificial light, especially blue light from screens, may increase the risk of breast cancer in addition to its negative effects on hormone balance, immunological function, and weight management ^8,9^. Nevertheless, while these factors may elevate the risk of breast cancer, the patterns of gene expression also vary from these factors.

Transcriptomics examines RNA transcripts, offering insights into gene expression patterns across many etiological causes^10,11^. Transcriptomic analysis is essential in breast cancer research for identifying gene expression profiles linked to breast cancer across various aetiologies. Previous investigation on transcriptomics has identified number of biomarkers that have advanced the diagnosis, prognosis, and treatment decision-making of breast cancer^12,13^. The pursuit of finding common essential genes while targeting novel indicators unique to induced environmental factors, such as medication and hormone imbalances, diet and nutrition, physical exercise, and nocturnal light exposure, remains still unexplored. Furthermore, the common genes identified in these etiological factors may significantly serve as biomarkers.

Biomarkers are crucial in contemporary medicine as they offer quantifiable indicators of biological processes, disease conditions, and therapeutic results. They are essential for precise diagnosis, prognosis, and therapy results^14,15^. In oncology, prevalent biomarkers such as carcinoembryonic antigen (CEA) and prostate-specific antigen (PSA) are employed for monitoring disease development, early detection, and cancer screening^15^. Prognostic indicators provide critical information regarding disease progression and the expected treatment trajectory for a patient. Elevated HER2 protein levels in HER2-positive breast tumours facilitate accelerated tumour progression. Treatment for HER2-positive breast cancer includes HER2-targeted therapies including trastuzumab and pertuzumab. Ki-67, a protein associated with cellular proliferation, is an indicator of rapid tumour progression and poor prognosis. Bowel and ovarian cancers have a higher incidence in those possessing BRCA1 or BRCA2 mutations ^16–21^. Hence, improving breast cancer treatment may be as simple as finding common indicators across several etiological factors. Finding the shared genes linked to different causes is, thus, believed to be beneficial. In addition, many etiological factors share genes that could be used as therapeutic targets.

Hence, we analyse gene expression datasets linked to three potential causes: hormonal and pharmaceutical imbalance, food and nutrition, and exercise and exposure to light at night. Our goal is to discover key genes that can be used to develop more effective biomarker-targeting approaches. Molecular mechanisms behind these common genes linked to cancer’s aetiology are also the subject of our investigation. Lastly, we use drug-complex analyses to validate these therapeutic targets through molecular simulation.

## Methods and Materials

### Selection of datasets

Sequencing datasets namely GSE33447, GSE244354, and GSE233242 were retrieved from National Centre for Biotechnology Information (NCBI) representing gene expression profile of stress, drug and hormonal imbalance, diet and nutrition, and physical exercise and light exposure at night mediated breast cancer samples from female respectively^22^. SRA-toolkit and fasterq-dump program were utilized to retrieve Sequence Read Achieve (SRA) data and their additional processing respectively^23^. Both of the tools can help in the collection of datasets from NCBI^24^.

### Data processing

#### Building genome index

The Homo sapiens reference genome GRCh38.p14 was obtained from the National Centre for Biotechnology Information (NCBI). Subsequently, the HISAT2 (Hierarchical Indexing for Spliced Alignment of Transcripts 2) tool was employed to create an index from the complete file containing the sequence of the human genome^25^. HISAT2 is a tool that can utilize the reference genome to a format that is well-suited for quick read alignment, guaranteeing accurate alignment of sequencing reads to the reference genome^26^.

### Trimming FASTQ Files

Trim Galore was used to remove unnecessary parts of the sequence from the raw FASTQ file including low-quality bases and adapter sequences to improve the accuracy for downstream analysis^27^. This tool provides quality trimming options to process raw FASTQ files.

### Aligning reads to genome

To align the trimmed sequence from upstream to the reference sequence, HISAT2 tool was used. This tool helps in identification of transcriptome characteristics and evaluate the genetic expression level of the samples under study. The output file is provided in SAM format ^26^.

### Converting SAM to BAM

For easier analysis in further processing steps and compatibility in storing the datasets, SAM (Sequence Alignment/Map) files are converted into BAM (Binary Alignment/Map) format by samtools^28^. The step significantly reduces file size and increases efficiency in storage^29^.

### Performing feature counting

Feature counting was done by utilizing ‘htseq-count’ which is a command of HTSeq that helped to determine the number of reads that overlapped with annotated genomic features^30^. High-Throughput Sequencing (HTSeq) is a tool that provides useful information about the level of gene expression and investigation of differential expression.

### Filtering count results

To eliminate any impurities and superfluous information or metadata in the datasets, the output from ‘htseq-count’ files was passed through the RNA-Seq analysis pipeline. This pipeline has the function of producing consistent results by using unprocessed sequence data into insightful data for rigorous study of study samples^30^.

### Identification of DEG genes

Using Cytoscape software^31^ (v3.869) and the InteractiVenn ^32^tool (http://www.interactivenn.net/), the identification of common dysregulated (both upregulated and downregulated) genes from the selected datasets was done^33^. Additionally, the resultant network displayed the genes that the datasets have in common.

### PPI network for common gene identification

Cytoscape and the STRING (Search Tool for the Retrieval of Interacting Genes/Proteins) ^34^(http://stringLdb.org/) database were leveraged to construct Protein-Protein Interaction (PPI) networks pertinent to breast cancer. With the help of search tools from STRING database, the interaction between the differentially expressed genes were identified which were analyzed in Cytoscape with a cutoff score of > 0.5^35^. Further, PPI networks were built and hub and bottleneck genes were investigated using the cytoHubba plug-in installed in the same software^36^.

### Gene ontology and pathway enrichment analysis

The DAVID (Database for Annotation, Visualization and Integrated Discovery) (https://david.ncifcrf.gov/) gene analysis tool was used to investigate breast cancer-related biological components^37^. This tool revealed molecular interaction and signalling cascade networks related to cell cycle regulation, hormone signalling, and DNA damage response in numerous subcellular sites and structures^38^.

### Protein and ligand preparation for molecular docking

The three-dimensional structures of selected downregulated and upregulated genes namely ZCCHC7 (PDB ID: Q8N3Z6) and UBE2J2 (PDB ID: Q8N2K1) were searched in RCSB PDB ^39^(https://www.rcsb.org/) for collecting their respective PDB file containing X-ray crystallographic coordination. The PDB files were prepared for docking by removing co-factors, water, and metal ions in BIOVIA Discovery Studio Visualizer Tool (v16.1.0.41) ^40^Further, Drug Bank was utilized for searching potential compounds with interaction capability with selected gene products and Avogadro tool^41^ for generating the ligand file suitable for molecular docking analysis.

### Molecular docking analyis

The molecular docking analysis was performed using PyRx virtual screening software^42^. Grid box parameters were applied to the generated crystal structures of ZCCHC7 (center_x = - 1.73, center_y = -5.36, center_z = 7.12, size_x = 114.3, size_y = 101.84, size_z = 99.80) and UBE2J2 (center_x = 16.47, center_y = 5.30, center_z = 2.72, size_x = 50.22, size_y = 32.75 size_z = 41.66) to analyze their interactions with selected ligands^43^. BIOVIA Discovery Studio Visualizer Tool (v16.1.0.41) was used to visualize the interactions.

### Molecular dynamics (MD) simulation

The GROMACS (GROningen MAchine for Chemical Simulations) (version 2020.6) program^44^ was utilized for molecular dynamic simulation for 100 ns. GROMOS96 43a1 force field was used to carry out the studies of root means square deviation (RMSD), root mean square fluctuation (RMSF), radius of gyration (Rg), and solvent accessible surface area (SASA) analysis. RMSD reflects the deviation in structural changes, RMSF searches flexibility or mobility of atoms over time, Rg values predict the folding nature of the overall structure, SASA represents the accessibility to any particular solvents. Lastly, ggplot2 in R was applied for generating plots for visualization^45^.

## Results

### Analysis of Gene Expression Profiles in Breast Cancer

Volcano plots illustrate the differentially expressed genes (DEGs) between tumor and normal samples in three distinct datasets containing gene expression profile of drug and hormonal imbalance, diet and nutrition, and physical exercise and light exposure at night in breast cancer patients. From the dataset of stress, drug and hormonal imbalance in breast cancer samples (patient and control), it was found that 1871 genes were upregulated and 823 genes were downregulated based on the magnitude of changes in gene expression (fold change) and their statistical significance (-log10 adjusted p-value) (**Supplementary Table 1**). On the same basis, diet and nutrition induced breast cancer showed 2538 upregulated and 2142 downregulated genes (**Supplementary Table 2**). Lastly, physical exercise and light exposure at night in breast cancer samples displayed 4853 upregulated and 6773 downregulated genes (**Supplementary Table 3**). Each dot on the plot representing a gene, and its position is dictated by its fold change (x-axis) and adjusted p-value (y-axis) was illustrated in **Figures 1A, 2A, 3A**.

**Figure 1:**
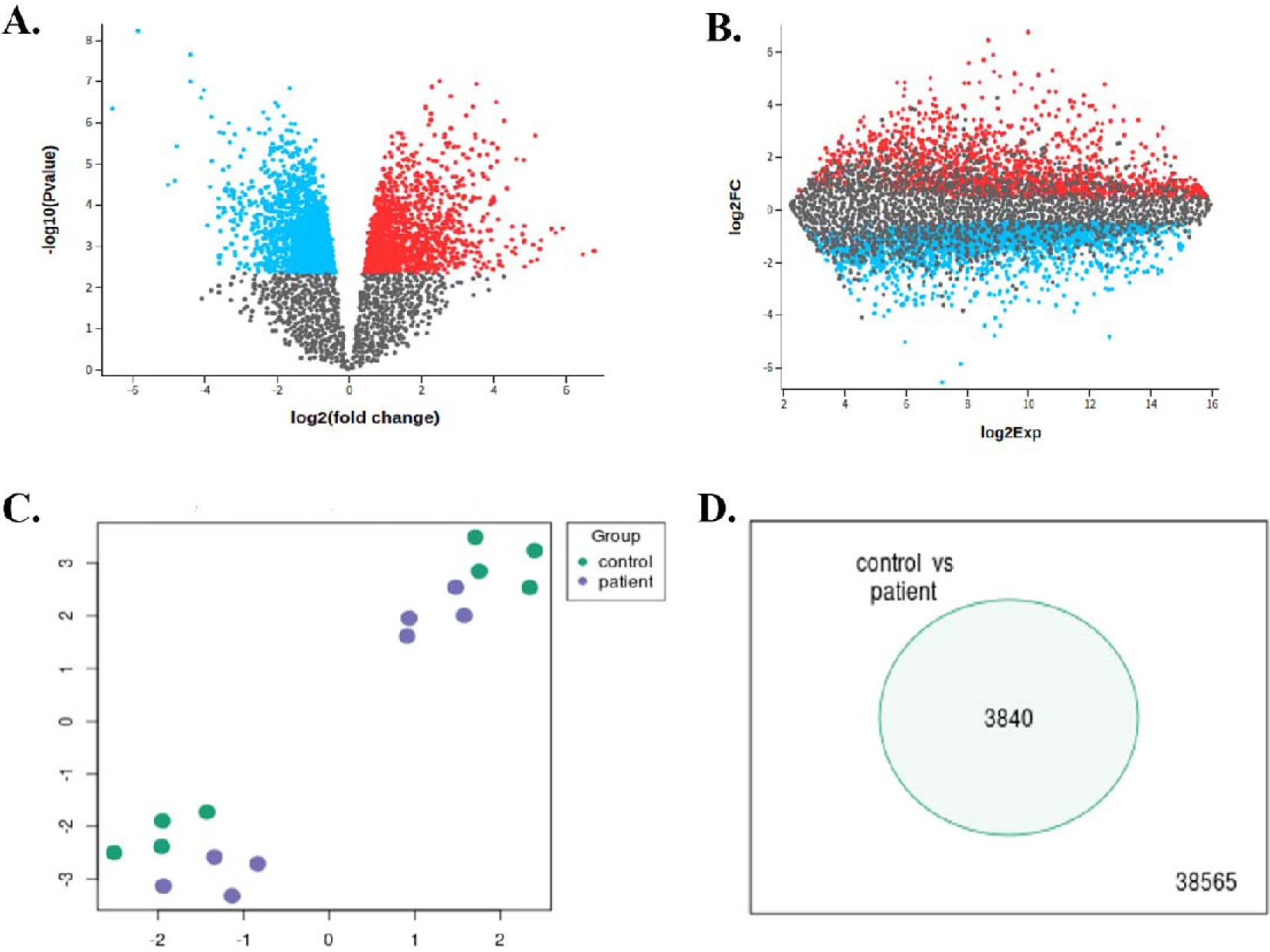
Gene expression profiles of Stress, Drug and Hormonal imbalance factor. (A) A volcano graphic shows the statistical significance (-log10 p-value) and gene expression change (log2 fold change). Red dots show upregulated genes in the patient cohort, blue dots downregulated genes, and grey dots unaffected genes. (B) MA plot: Shows log2Exp on the x-axis and log2FC on the y-axis. This graph shows the relationship between gene expression levels and group variation. Blue dots indicate downregulated genes, red dots indicate upregulated genes, and grey dots indicate no change. C) Scatter plot: Shows the expression levels of a selected selection of genes in the control (green dots) and sick (purple dots) groups. This graphic shows gene expression differences. D) Venn diagram: Shows the number of genes expressed solely in the control group and shared by both groups. This graph compares gene expression profiles of the two groups.

**Figure 2:**
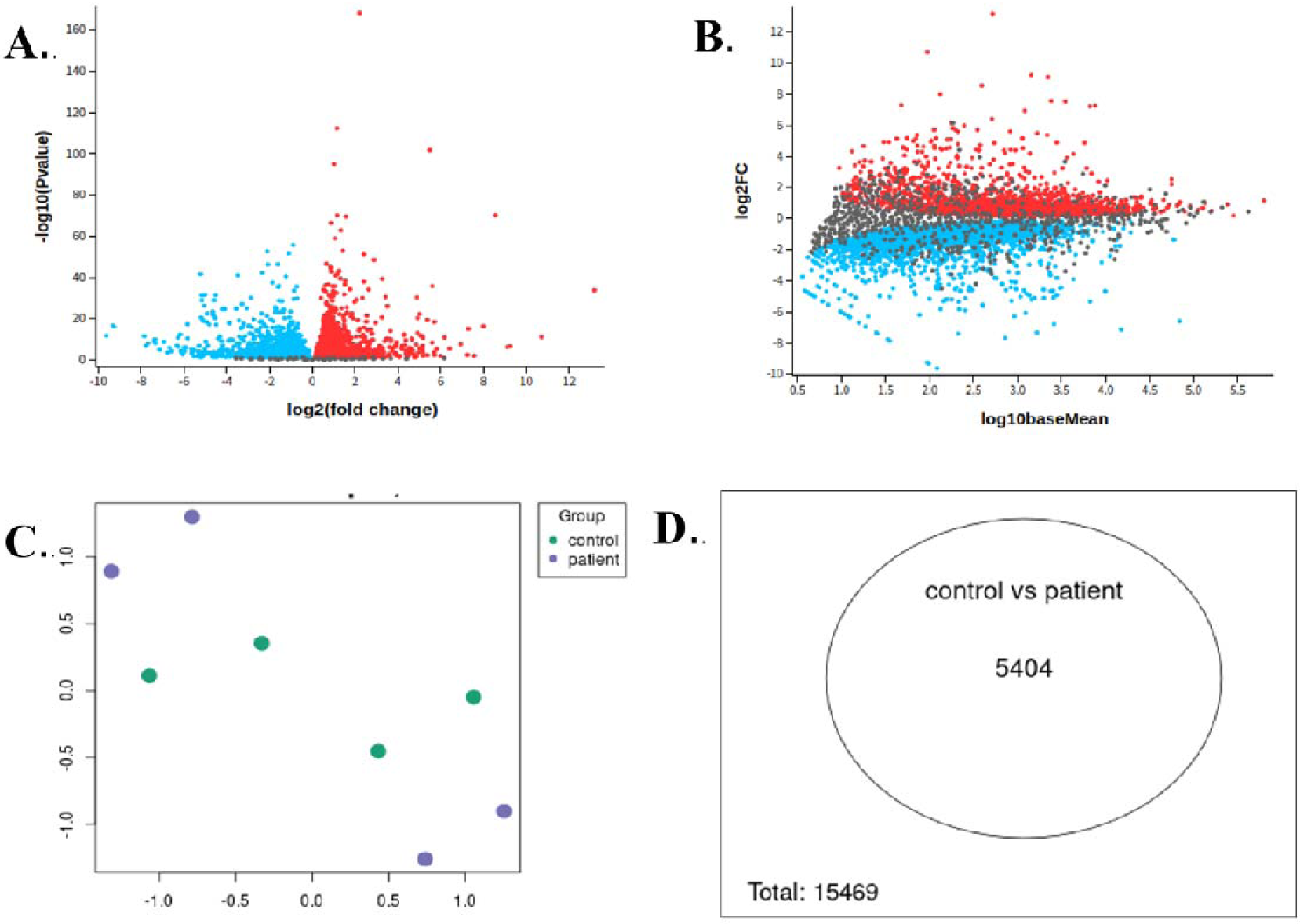
Gene expression profile of diet and nutrition-induced breast cancer. (A) Volcano plot (B) Mean-difference plot (C) UMAP plot (D) Venn diagram. The downregulated genes are specified by blue dots, red dots indicate upregulated genes and the genes that are not significant are indicated by grey dots.

**Figure 3:**
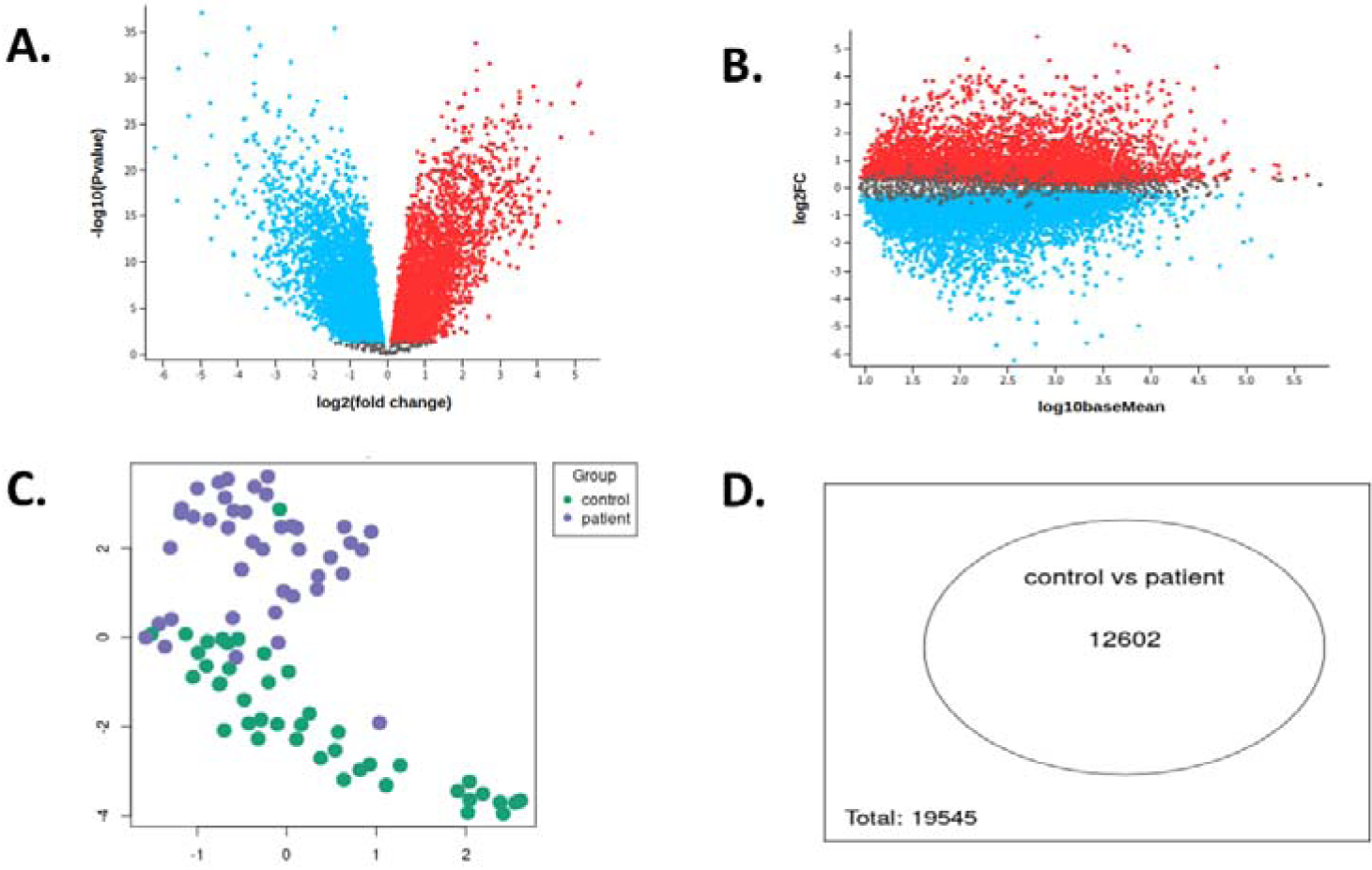
Gene expression profile of physical exercise and light exposure at night. (A) Volcano plot (B) Mean-difference plot (C) UMAP plot (D) Venn diagram. The downregulated genes are specified by blue dots, red dots indicate upregulated genes and the genes that are not significant are indicated by grey dots.

Mean-Difference plots, which illustrate each gene’s average expression level against its log2 fold change, offer an additional perspective on gene expression patterns. Finding genes with significant expression differences between tumor and normal samples is made easier with the help of this visualization. The terms "upregulated" and "downregulated" refer to genes that are situated above or below the horizontal line (which indicates no fold change). Genes that exhibit significant changes in expression, either upregulated or downregulated, were positioned away from the center, suggesting a possible role for these genes in the development of breast cancer (**Figures 1B, 2B, 3B**). The steeper the slope of the line connecting a gene to the origin, the greater the expression difference between tumor and normal samples.

Datasets of stress, drug and hormonal imbalance in breast cancer samples included 8 patients and 8 controls. Additionally, diet and nutrition induced breast cancer datasets had 4 patient and control samples for each category. Likewise, physical exercise and light exposure at night in breast cancer datasets contained 43 samples for both patient and control.

Uniform Manifold Approximation and Projection (UMAP) plots show gene expression patterns connecting samples with breast cancer in tumour and normal samples and reduce dimensionality (**Figures 1C, 2C, and 3C**).

Venn diagrams demonstrated 3840, 5404 and 12602 genes were dysregulated in datasets of stress, drug and hormonal imbalance, diet and nutrition, and physical exercise and light exposure at night in breast cancer samples among the total of 38565, 15469, 19545 genes respectively (**Figures 1D, 2D, 3D**). By identifying both common and distinct DEGs through overlapping and non-overlapping regions, this approach sheds light on similar molecular pathways and variations particular to the dataset.

### Upregulated and downregulated gene analysis revealed common dysregulated genes

From the datasets, the common up- and down-regulated genes of breast cancer were identified using the Cytoscape software v3.8 and the InteractiVenn tool (http://www.interactivenn.net/) (**Supplementary Table 4**). There were twenty-two upregulated genes and four downregulated genes among the datasets. DESeq2 tool in R was used to create feature counts from RNA sequencing data processing pipeline for visual analysis (**Figure 4**).

**Figure 4:**
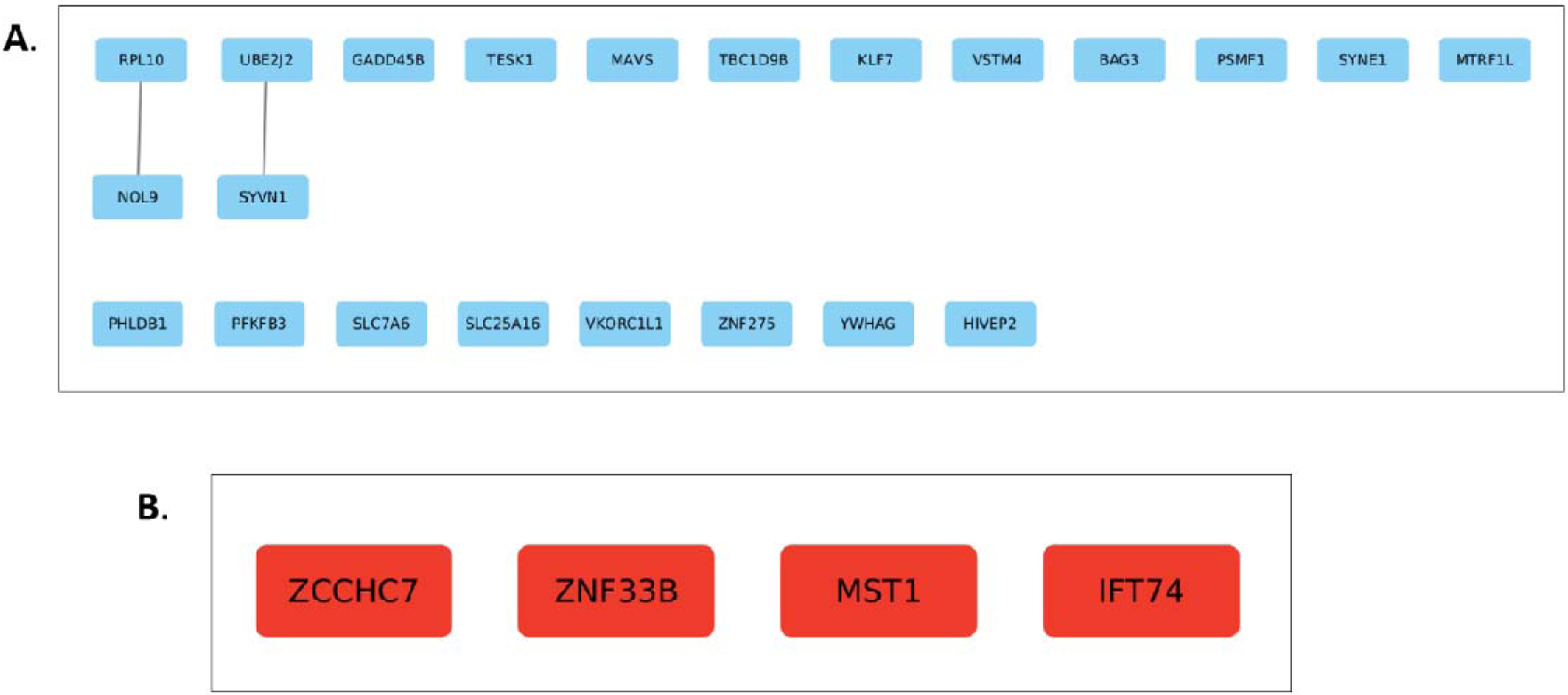
Identification of the common dysregulated genes among all the etiologic factors. (A) upregulated genes (B) downregulated genes.

Findings from this investigation indicated that 4 genes were downregulated (ZCHHC7, ZNF33B, MST1, IFT74) and 22 upregulated genes (PFKFB3, BAG3, HIVEP2, KLF7, TBC1D9B, VSTM4, GADD45B, MAVS, MTRF1L, NOL9, PHLDB1, PSMF1, RPL10, SLC25A16, SLC7A6, SYNE1, SYVN1, TESK1, YWHAG, UBE2J2, VKORC1L1, ZNF275), which may indicate that these genes are involved in key processes that lead to the development of breast cancer. Since these common DEGs consistently exhibit dysregulation across a variety of datasets, they may offer attractive targets for therapy.

### Functional enrichment confirms the dysregulated genes’ involvement in important function

Gene Ontology (GO) provided insights into the biological processes, cellular components, and molecular functions associated with the identified DEGs (**Figures 5 and 6**). Transcriptional regulation, ubiquitin-dependent protein degradation, and ATP binding were among the activities for which upregulated genes were specifically enriched. These results imply that changes to these mechanisms could facilitate the development of breast cancer. For example, changes in ubiquitin-dependent protein degradation may disrupt protein homeostasis, and disruption of transcriptional control can result in abnormal patterns of gene expression, both of which can aid in the development of cancer.

**Figure 5:**
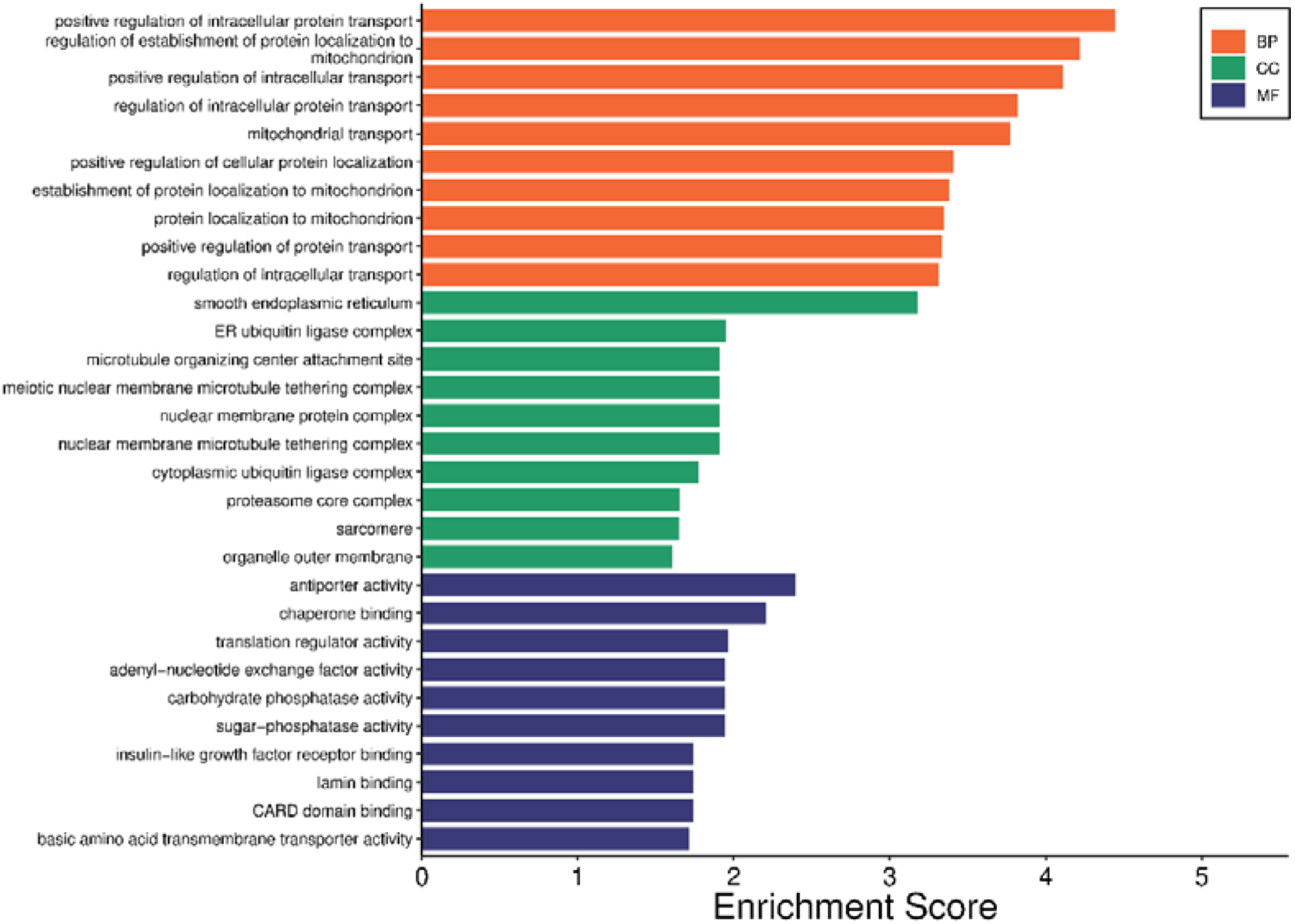
Gene ontology of upregulated genes. The bar chart displays the enrichment scores for GO elements that were significantly enriched. The three separate ontologies from which the values are obtained include Biological Process (CC, green), Molecular Function (MF, blue), and Biological Process (BP, orange). The x-axis of the graph shows the enrichment score, which indicates how over-represented a GO term is in the gene collection. A stronger correlation between the gene set and its matching GO term is indicated by higher scores.

**Figure 6:**
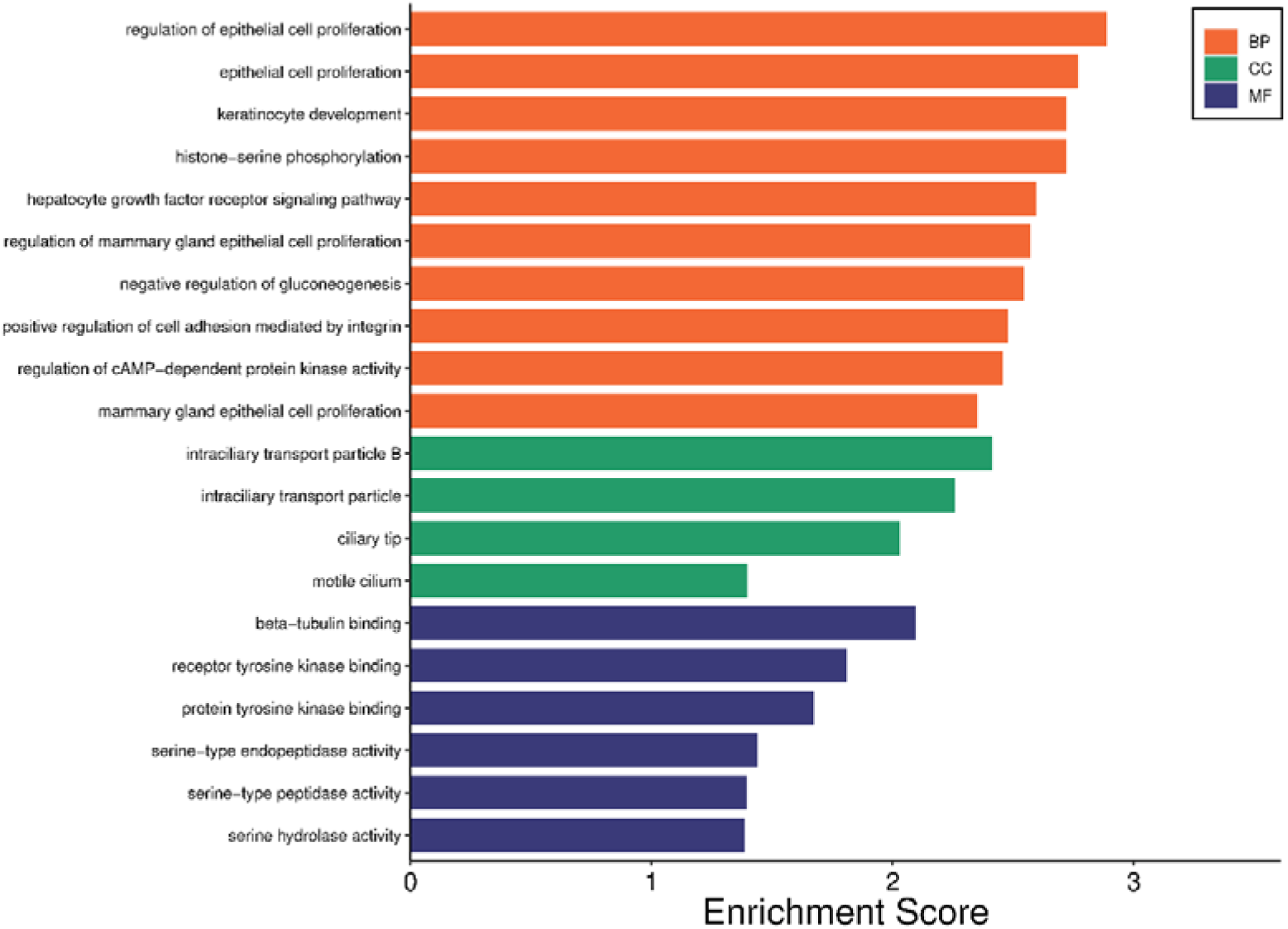
Gene ontology of downregulated genes. The three distinct ontologies from which the values are derived are Cellular Component (CC, green), Molecular Function (MF, blue), and Biological Process (BP, orange). The higher the enrichment score, the higher the stronger relation of the genes with the GO terminology.

Upregulated genes were substantially enriched in pathways linked to the cell cycle, endoplasmic reticulum (ER) stress, and responses to viral infection, according to KEGG pathway analysis (**Figures 7 and 8**). These findings suggest that the deregulation of these pathways plays a role in the etiology of breast cancer. For instance, uncontrolled proliferation of cells can result from cell cycle dysregulation, and autophagy or apoptosis can be triggered by ER stress, both of which may progress cancer.

**Figure 7:**
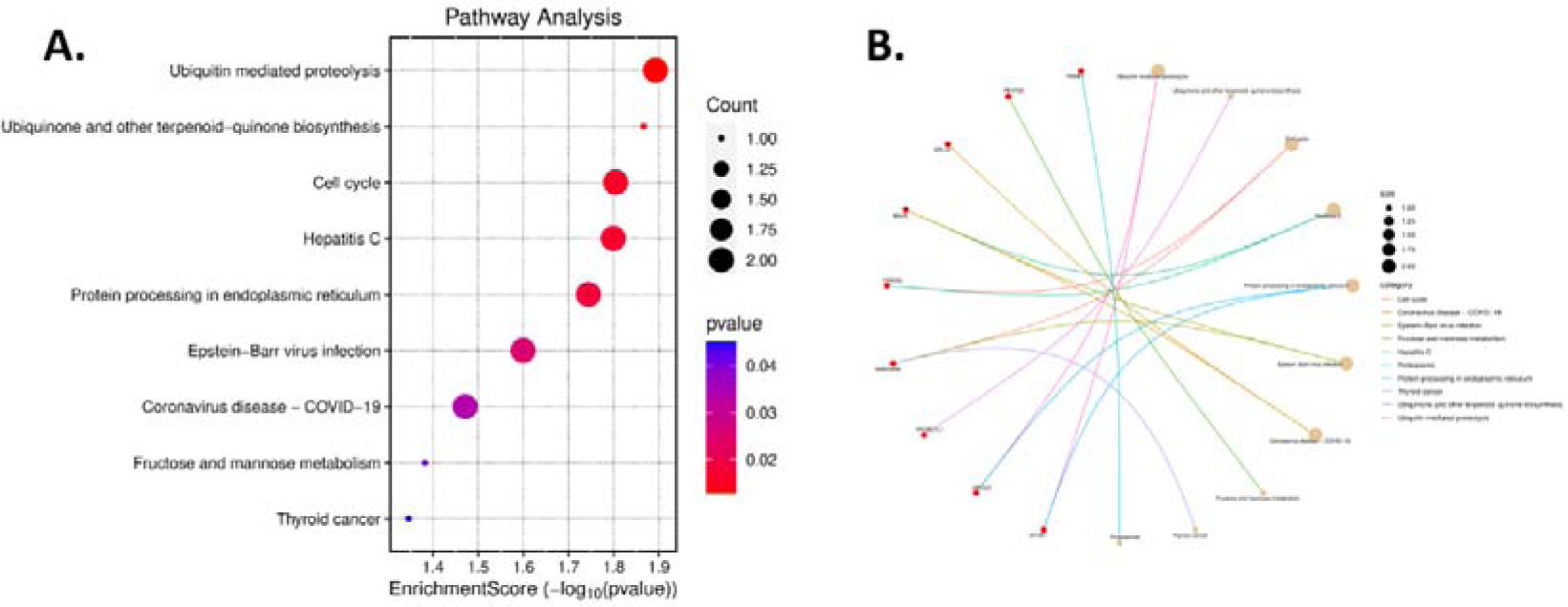
Pathway of upregulated genes. (A) enrichment score dot plot: shows the enriched routes on the x-axis, with their enrichment scores (-log10 p-values) used to sort them. Dot size represents the number of genes associated with each pathway, and colour intensity represents the adjusted p-value. (B) cnet plot: shows how genes are linked to enriched pathways graphically. Genes (circles) and routes (squares) are shown as nodes, and the relationships between them are shown by lines. Each pathway category is denoted by a different edge colour.

**Figure 8:**
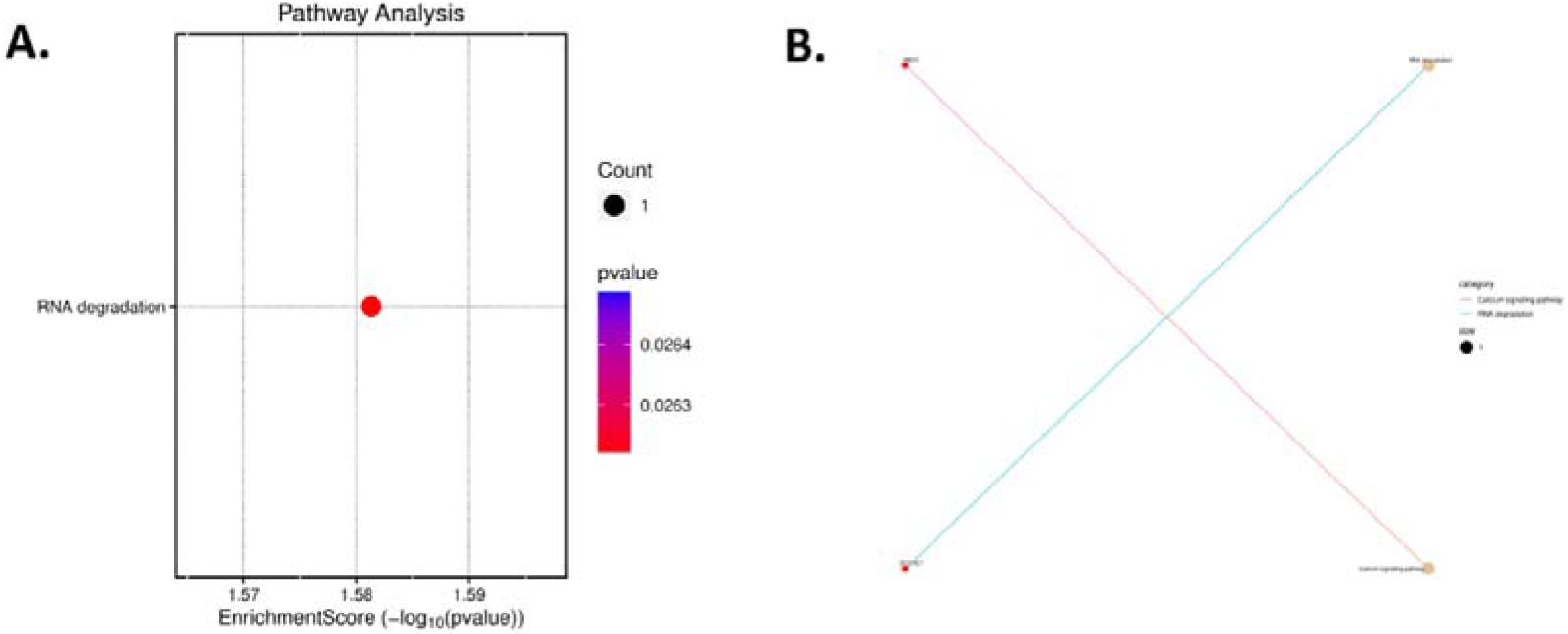
Pathway enrichment of downregulated genes. (A) enrichment score dot plot: displays enriched routes on the x-axis by enrichment scores (-log10 p-values). Dot size indicates the number of genes per route, and colour intensity indicates the adjusted p-value. (B) cnet plot: shows gene-enriched pathway linkages. Nodes are genes (circles) and pathways (squares), while edges (lines) show their interactions. Edge colour matches pathway category.

### Hub and Bottleneck Genes confer the central influence on the interactions

Hub and bottleneck genes, which are essential for network connectivity and information flow, were found by PPI network analysis (**Figures 9 and 10**). Hub genes identified from the upregulated genes on three etiological factors are RPS14, JUN, MYC, UBC, UBB, IL6, UBA52, RPS18, RPS27A, EGFR, GRB2, PTPRC, FCGR3A, CASP3, STAT1, KRAS, H3C12, TGFB1, CD19, CD74, RPS27A, RPS2, RPL5, RPL4, RPS14, ACTB, UBC, UBB, RPS16, RPS6. On the other hand, bottleneck genes identified from the upregulated genes are RPS6, RHOA, MYC, PPARGC1A, UBC, ACO2, ALB, FOS, EGFR, RPS27A, GRB2, CTSD, RAB8A, MDM2, STAT1, P4HB, KRAS, H3C12, CXCR4, CD74, RPS27A, RPL8, ACO2, SRSF1, CTNNB1, CCT7, HSP90AA1, ACTB, UBC, RPS6. Hub genes identified from the downregulated genes on three etiological factors are ESR1, GAPDH, HSP90AA1, HSPA4, POLR2B, TRIM28, HSP90AB1, EP300, HDAC1, H2BC21, ACTB, CTNNB1, HSPA8, ATM, EZH2, AKT1, BRCA1, SRC, RAD51, RPL19, H2AC20, ISG15, H2BC5, H2BC21, GSK3B, CDKN2A, SIN3A, TP53, CHEK2, RIGI, PINK1, H2BC12, ATM, LAMP1, TBK1, H2AC18, NOTCH1, ULK1, RB1, SQSTM1, IRF1, RPL7, RPL15, RPL18A, RPS4Y2, RPS3, RPL23, RPS14, RACK1, RPL22, RPS9, RSL24D1, RPS29, RPL35, RPS10, RPL21, RPSA, RPL39, RPL13, EEF1B2. Further, bottleneck genes identified from the downregulated genes are CCNA1, DTNA, HIPK3, XPO1, CDH1, CTNNA1, UBB, SRPK1, CTNNB1, HSP90AB1, APC, IGF1R, SNCA, IFT46, GSK3B, PACSIN1, TOP1, TRIM28, KRAS, EP300, CAV3, TP53, LGALS3, HSPA1A, SREBF1, POLR1B, PLCG1, SOS1, ISG15, IGF1, DMD, TP73, ZNF2, CTSD, KCNH2, LRRK2, RAB8A, SRC, NCL, AHSG, CYHR1, HNRNPDL, RACK1, RGN, ACTB, EHD1, GNG11, SNAP29, SRSF1, ADCY4. Because these genes play a major role in network dynamics, they may be interesting targets for therapy. In the network, hub genes are highly connected to other genes such as SYVN1, UBE2J2 and KLF7 whereas bottleneck genes such as ZNF33B, IFT74 and ZCCHC7 serve as vital connectors between other network modules. By focusing on these genes, it may be possible to interfere with the network and inhibit the spread of cancer.

**Figure 9:**
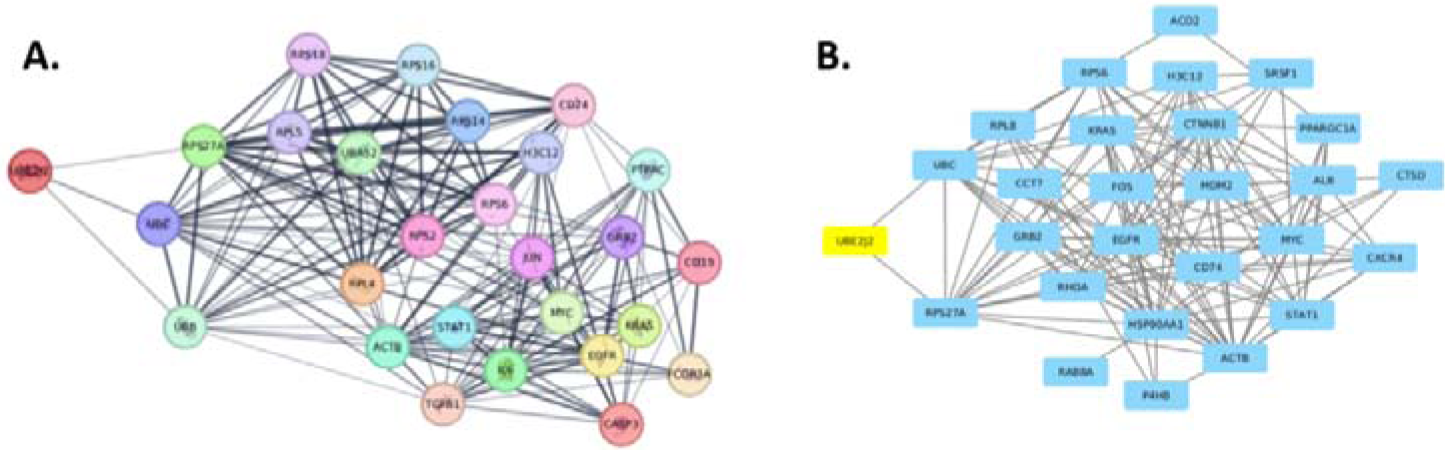
The interactome of UBE2J2 target. A) hub genes of UBE2J2 target and B) bottleneck genes of the target.

**Figure 10:**
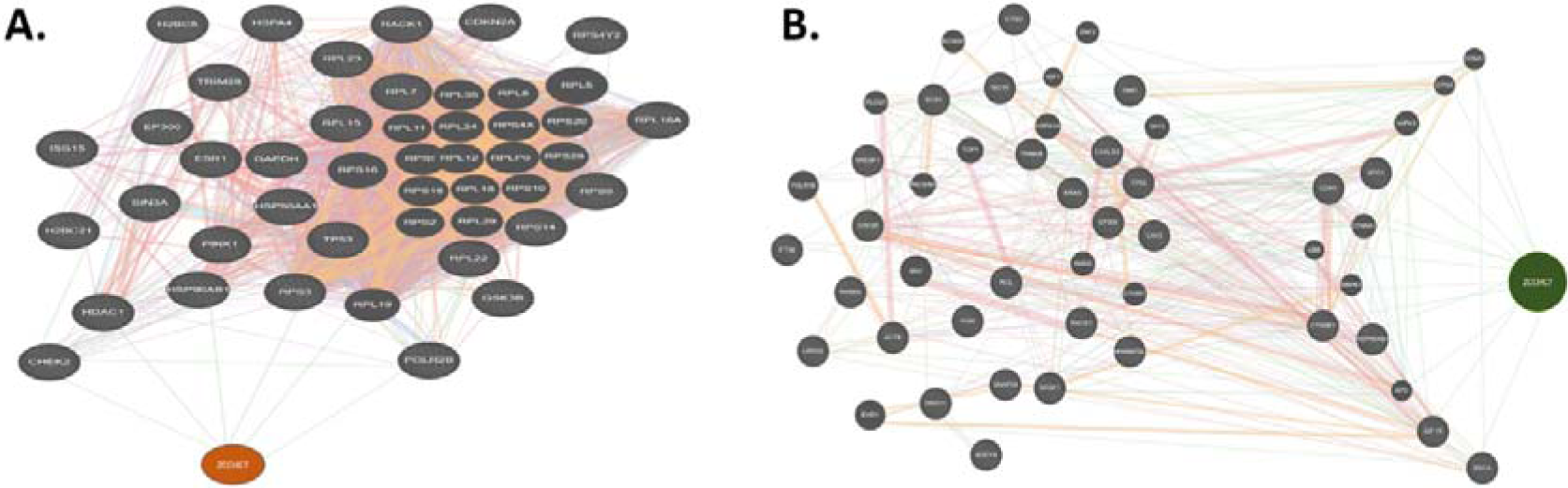
The interactome of ZCCHC7 target. A) Identification of partner genes of ZCCHC7 and B) bottleneck genes of the target ZCCHC7.

### Drug-Target Interaction Prediction confirms their target-like nature

Based on the computational study, two possible drug-like substances that may interact with hub and bottleneck genes (UBE2J2 and ZCCHC7) are Artenimol and DB07679. (**Figure 11**). The magnitude of these connections was determined by binding affinity calculations, which also suggested possible therapeutic effectiveness. While DB07679 is an unidentified investigational chemical, Artenimol is an antimalarial medication with possible anticancer action. According to **Tables 1 and 2**, these compounds have the ability to modify the activity of ZCCHC7 and UBE2J2, respectively, based on their calculated binding affinities.

**Figure 11:**
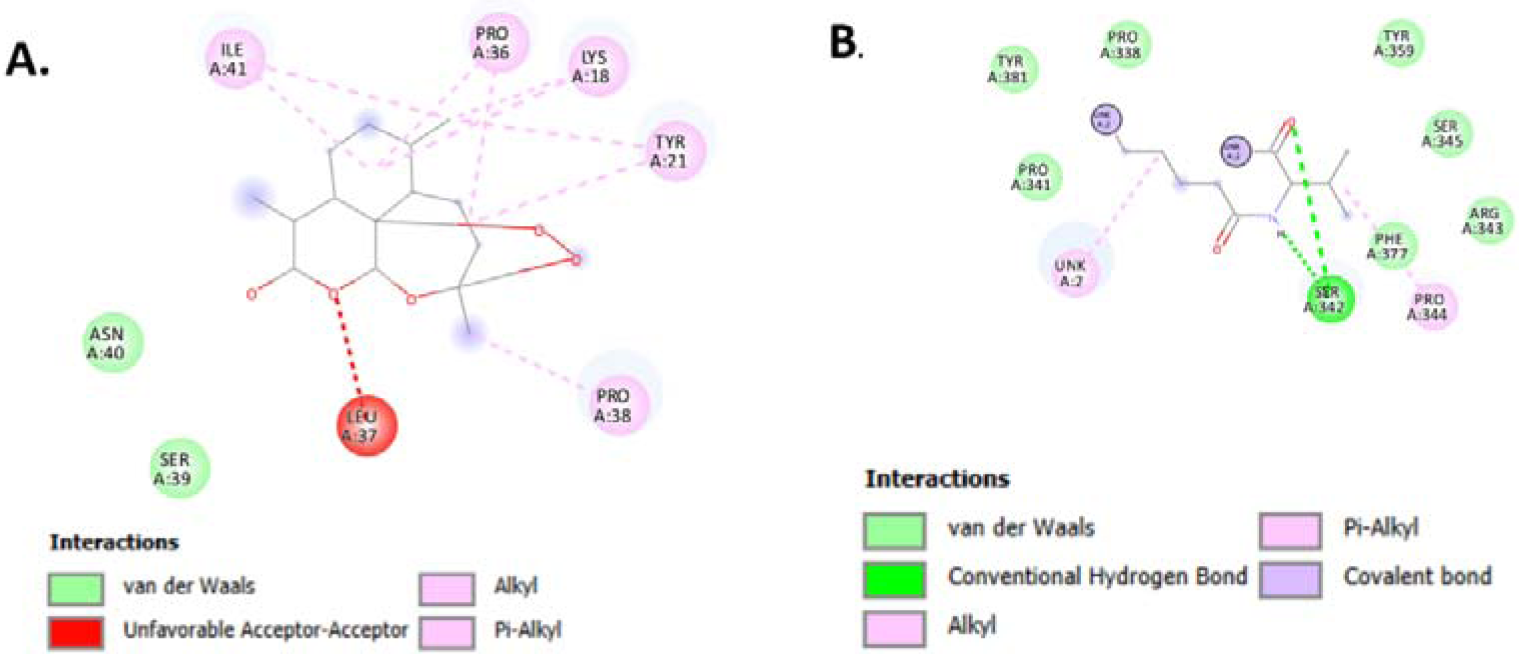
2D interaction between (A) UBE2J2-Artenimol (receptor-ligand) complex, (B) ZCCHC7-DB07679 (receptor-ligand) complex.

**Table 1:** List of hub and bottleneck genes.

| Upregulated gene |  |  | Downregulated gene |  |  |
| --- | --- | --- | --- | --- | --- |
| Symbol | Hub gene | Bottleneck gene | Symbol | Hub gene | Bottleneck gene |
| SYVN1 | UBB, UBC, RPS27A | UBC, RPS27A, P4HB, HSP90AA1 | ZNF33B | TRIM28 | TRIM28 |
| UBE2J2 | UBB, UBC, RPS27A | UBC, RPS27A | IFT74 | - | IFT46 |
| KLF7 | - | - | ZCCHC7 | CHEK2, HDAC1, HSP90AB1, RPS3, RPL19, POLR2B | CCNA1, DTNA, HIPK3, XPO1, CDH1, CTNNA1, UBB, SRPK1, CTNNB1, HSP90AB1, APC, IGF1R, SNCA |

**Table 2.**
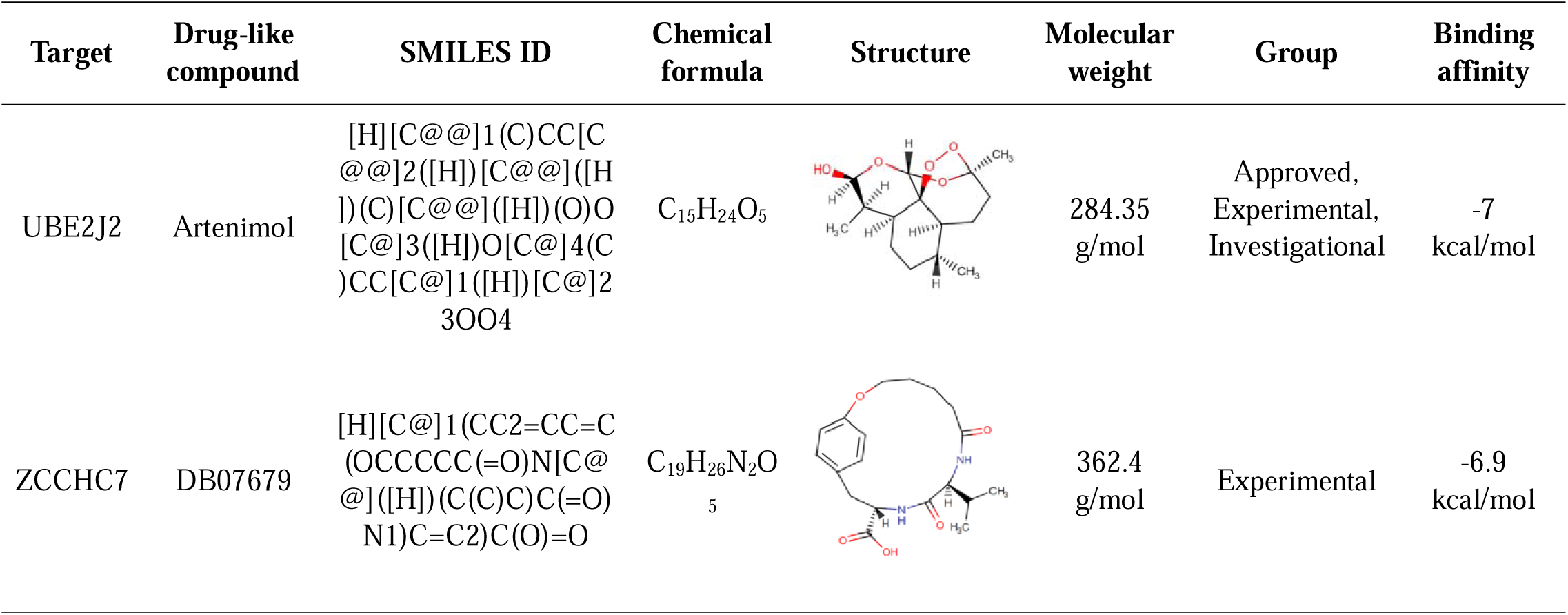
Selected drug-like compounds and their properties.

Molecular dynamics simulations assessed the stability and dynamic behavior of the predicted drug-target complexes (**Figures 12 and 13**). The receptor-ligand interaction of UBE2J2-Artenimol had some fluctuations in RMSF, RMSD, Rg and SASA analysis. For RMSF, the highest peak was between residue 120 and 140. On the other hand, the interaction of ZCCHC7-DB07679 had relatively no fluctuations in all four metrices. In RMSF, the value was below 2.5 for 100 ns time. However, RMSD showed upward trends and was relatively stable after 30 ns. However, the Rg of this interaction showed the opposite trend with downward values over time which slightly increased after 80 ns. Further, SASA results showed some fluctuating nature from the beginning to the end over time. Overall, the simulations demonstrated the stability of the drug-target complexes over time, as well as the possibility for the medications to modify the activity of the corresponding targets.

**Figure 12:**
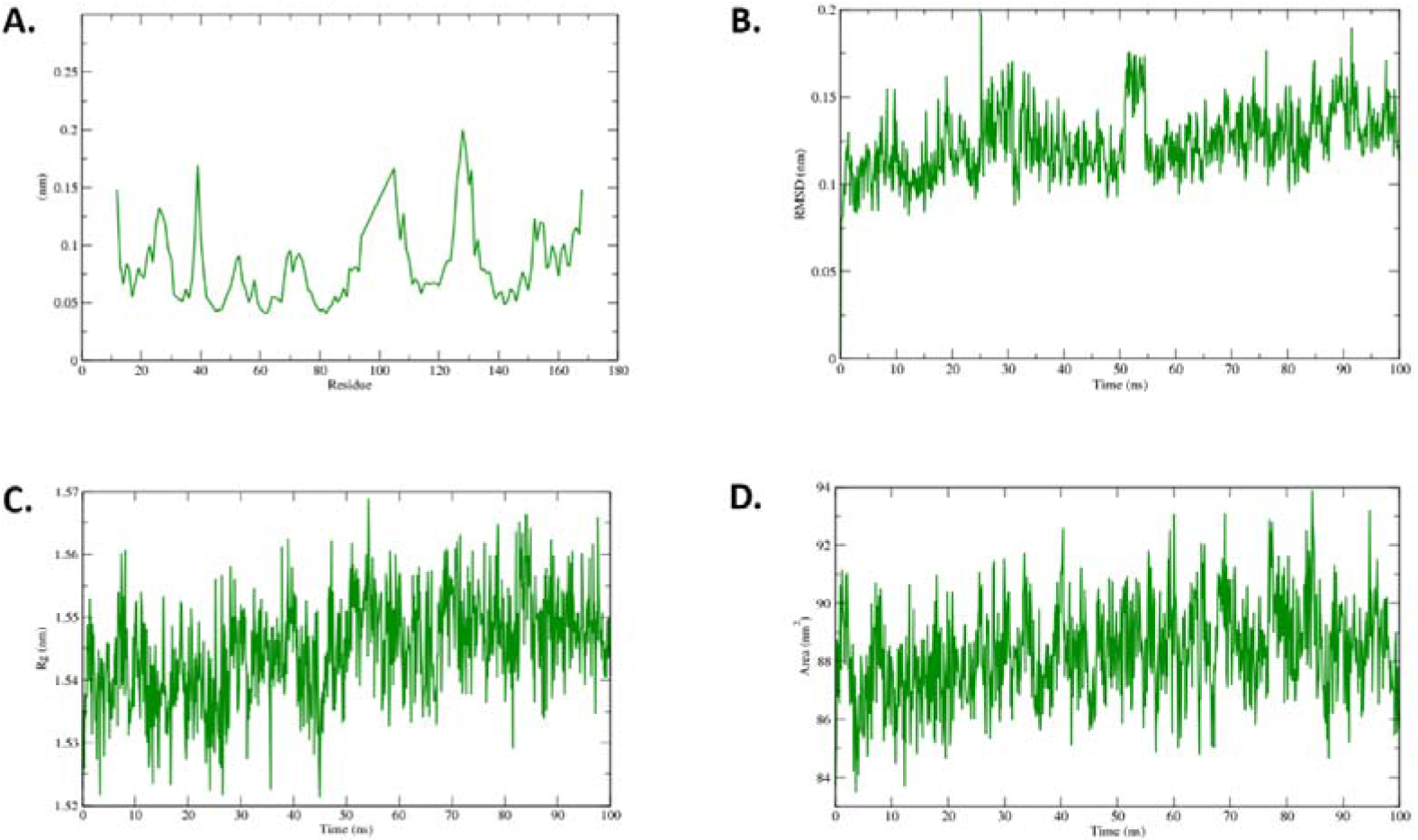
Molecular dynamic simulation of UBE2J2-Artenimol (receptor-ligand) complex. (A) RMSF (B) RMSD (C) RG (D) SASA

**Figure 13:**
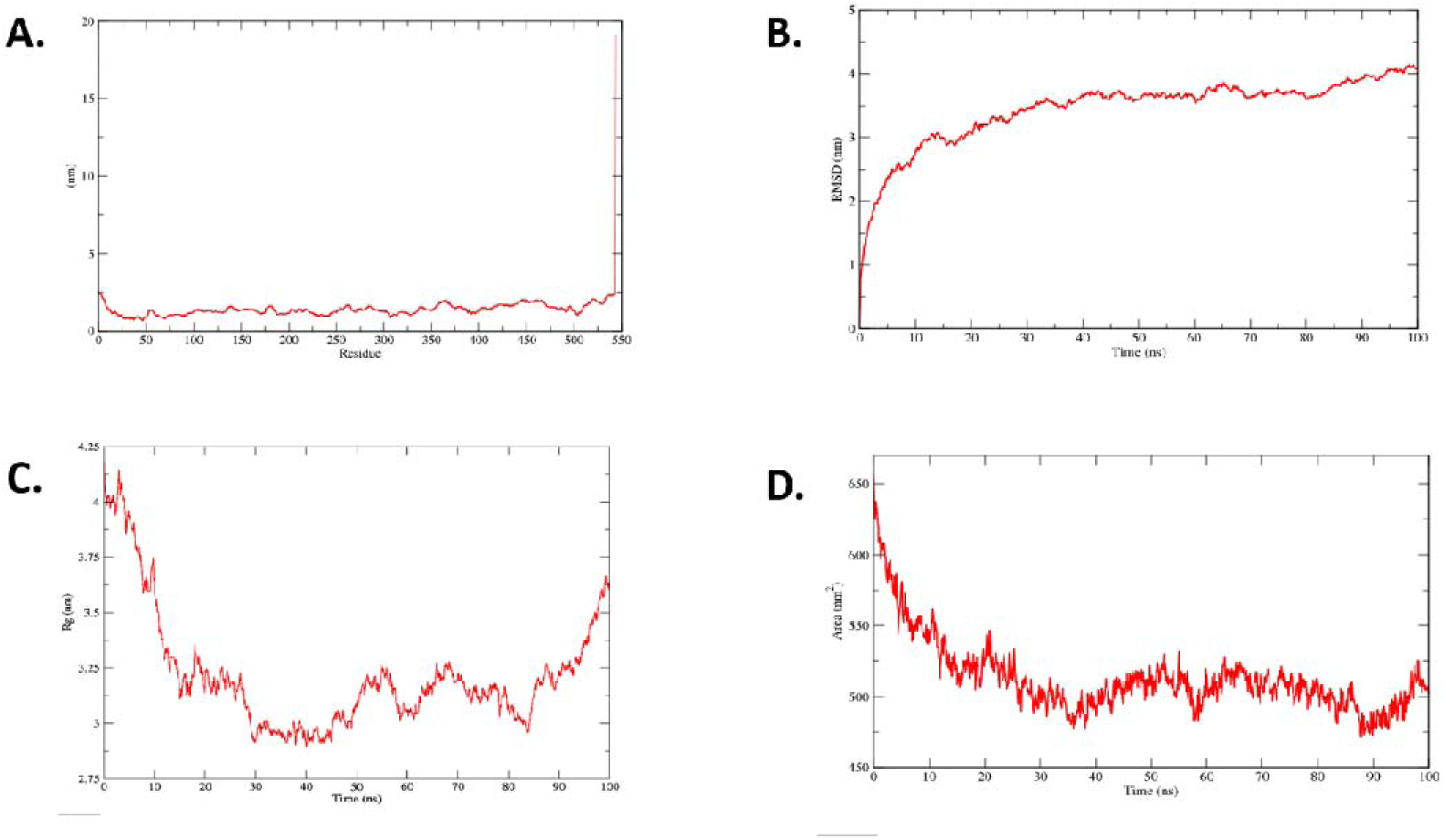
Molecular dynamic simulation of ZCCHC7-DB07679 (receptor-ligand) complex. (A) RMSF (B) RMSD (C) RG (D) SASA

## Discussion

Our study utilizes the datasets of gene expression profiles from 110 breast cancer study samples, which represent the etiological factors, stress, drug and hormonal imbalance (GSE33447), diet and nutrition (GSE244354), and physical activity and light exposure at night (GSE233242). Analysis of these datasets provides comprehensive insights for development of breast cancer and identifies potential therapeutic targets.

The analysis with different RNA-sequencing tools has shown that the genes that are responsible for breast cancer are dysregulated into the many etiologic factors. The analysis using Volcano, Mean-Difference, and UMAP plots illustrates genes that are expressed differently in malignant samples compared to normal samples (**Figures 1-3, Supplementary Tables 1-3**). This could be a result of the influence of a variety of factors that are associated with breast cancer.

Strikingly, Venn diagram confirms the common differentially expressed genes regardless of the etiologic factors **(Figures 1-3)**. In this connection, the InteractiVenn program and the Cytoscape software identify 22 shared upregulated genes and 4 shared downregulated genes which are consistent in their dysregulation patterns irrespective of etiological backgrounds **(Figure 4 and Supplementary Table 4)**. This observation aligns with previous research demonstrating the involvement of common genes and pathways in breast cancer pathogenesis, regardless of the specific risk factors^46,47^.

Additionally, it is highly anticipated to check the involvement of the identified genes in important functions which might contribute to the potential harmful impacts resulting from the breast cancer. Hence, functional enrichment analysis provides details about biological processes, cellular components, and molecular functions to suggest the impact of alterations associated with the identified genes in the previous steps **(Figures 5, 6 and Supplementary Tables 5-7, 8-10)**. Moreover, enrichment of upregulated genes in transcriptional regulation, ubiquitin-dependent protein degradation, and ATP binding processes shows the possibility of disrupting homeostasis and further leading to cancer^48,49^. The results are further supported by the KEGG pathway analysis that predicts the functional enrichment of pathways related to cell cycle, endoplasmic reticulum (ER) stress, and viral infection responses **(Figures 7, 8 and Supplementary Tables 11-12, 13-14)**. Impact of cell cycle dysregulation in uncontrolled proliferation, ER stress in autophagy and viral infection or chronic inflammation can be important factors in cancer development and progression^50–53^.

Protein-Protein Interaction (PPI) networks often represent the connection and interaction patterns of one protein with another. In this case of breast cancer, the assessment of identified dysregulated genes by PPI networks confirms hub and bottleneck genes namely UBE2J2 and ZCCHC7 **(Figures 9, 10 and Table 1)**. It was discovered that the interactome of upregulated genes had the most interacting partners with SYVN1 and UBE2J2, while the interactome of downregulated genes had the most connections with the ZCCHC7 gene. The development of breast cancer is influenced by each of these identified genes **(Figure 7-10)**. Recognizing their critical involvement in overall information flow in breast cancer development^54,55^, computational analysis is performed for therapeutic interaction analysis. UBE2J2 demonstrated druggable potential in docking, as existing medicines targeting this protein are present in the database, unlike those for the SYVN1 target **(Table 2).** Two drug-like compounds (Artenimol and DB07679) out of other compounds with connectivity to selected hub and bottleneck genes **(Supplementary Tables 15-18)** suggest potential therapeutic efficacy due to strong binding affinity (-7 kcal/mol and -6.7 kcal/mol respectively) illustrated by the molecular docking analysis **(Figure 11 and Table 2).** In addition to this, they demonstrated a variety of amino acids that interact with the drug Artenimol, such as lysine-18, tyrosine-21, proline-36, leucine-37, proline-38, serine-39, asparagine-40, and isoleucine-41 which had pi bonds, van der Waals interactions and other types of bonding. On the other hand, DB07679 also showed the interactions with proline-338, proline-341, serine-342, arginine-343, proline-344, serine-345, tyrosine-359, tyrosine-381 which had van der Waals, conventional hydrogen bonds and alkyl bonds **(Figure 11).**

Subsequent molecular dynamics analysis of these drug-target interactions indicates different metrics including RMSD, RMSF, Rg and SASA of the interactions **(Figures 12 and 13).** The dynamic behavior over the range of 100 ns is predicted which is expected to be highly reproducible under wet lab settings further strengthens the therapeutic importance of the selected drugs under breast cancer conditions. The outcomes from our investigation confirmed that the two targets could serve as prospective therapeutic targets, as similar studies also validated these results^33,56,57^.

Our study expands upon previous work by identifying hub and bottleneck genes and offers crucial information on differentially expressed genes in the presence of specified etiological variables. Hub and bottleneck genes confirm that UBE2J2 and ZCCHC7 interact with many genes involved in breast cancer events, and these two dysregulated genes are present in all etiologic factors. Thus, when these etiologic factors are present, these two genes UBE2J2 and ZCCHC7 are the common occurrences of breast cancer. We therefore hypothesise that the two genes found in this study may serve as a biomarker for the prognosis of breast cancer. Additionally, the drug binding analysis and molecular dynamics studies provide more evidence of their importance in breast cancer therapies aimed at reversing the disease’s course.

## Conclusion

In conclusion, the RNA-seq data, together with PPI and simulation approaches, indicated common dysregulated genes irrespective of etiological factors. Furthermore, we found two genes, UBE2J2 and ZCCHC7, that directly influence breast cancer development, as demonstrated by biological, molecular, and pathway enrichment analyses, indicating that these genes may serve as novel biomarkers. The investigation of drug binding and stability indicated their therapeutic potential for breast cancer treatment. Hence, we anticipate that findings of this study will significantly enhance the progression of breast cancer research towards the identification of novel biomarkers.

## Author Contributions

**Mohammad Uzzal Hossain:** Data curation, Formal analysis, Conceptualization, Methodology, Software, Visualization, Writing – original draft. **Mehak Ahmed:** Data curation, Formal analysis, Conceptualization, Methodology, Software, Visualization, Writing – original draft. **SM Sajid Hasan:** Data curation, Formal analysis, Methodology, Software, Visualization, Writing – original draft. **Mohammad Nazmus Sakib:** Data curation, Formal analysis, Software, Visualization, Writing – original draft. **A.B.Z. Naimur Rahman:** Data curation, Methodology, Software, Visualization, Writing – original draft. **Arittra Bhattacharjee:** Data curation, Validation, Visualization, Writing – review & editing. **Zeshan Mahmud Chowdhury:** Data curation, Validation, Writing – review & editing. **Ishtiaque Ahammad:** Data curation, Validation, Writing – review & editing. **Md. Mehadi Hasan Sohag:** Data curation, Validation, Supervision, Writing – review & editing. **Keshob Chandra Das:** Data curation, Validation, Writing – review & editing. **Md. Salimullah:** Conceptualization, Investigation, Resources, Supervision, Writing – review & editing, Data curation, Formal analysis.

## Acknowledgments

The authors have nothing to report.

## Data Availability Statement

All data generated or analyzed during this study are included in this published article.

